# The diazepam binding inhibitor’s modulation of the GABA-A receptor is subunit-dependent

**DOI:** 10.1101/2021.08.13.456294

**Authors:** Jennifer S. Borchardt, Lucas M. Blecker, Anton Tung, Kenneth A. Satyshur, Cynthia Czajkowski

## Abstract

First synthesized in the 1950s, benzodiazepines are widely prescribed drugs that exert their anxiolytic, sedative and anticonvulsant actions by binding to GABA-A receptors, the main inhibitory ligand-gated ion channel in the brain. Scientists have long theorized that there exists an endogenous benzodiazepine, or endozepine, in the brain. While there is indirect evidence suggesting a peptide, the diazepam binding inhibitor, is capable of modulating the GABA-A receptor, direct evidence of the modulatory effects of the diazepam binding inhibitor is limited.

Here we take a reductionist approach to understand how purified diazepam binding inhibitor interacts with and affects GABA-A receptor activity. We used two-electrode voltage clamp electrophysiology to study how the effects of diazepam binding inhibitor vary with GABA-A receptor subunit composition, and found that GABA-evoked currents from α3-containing GABA-A receptors are weakly inhibited by the diazepam binding inhibitor, while currents from α5-containing receptors are positively modulated. We also used *in silico* protein-protein docking to visualize potential diazepam binding inhibitor/GABA-A receptor interactions that revealed diazepam binding inhibitor bound at the benzodiazepine α/γ binding site interface, which provides a structural framework for understanding diazepam binding inhibitor effects on GABA-A receptors. Our results provide novel insights into mechanisms underlying how the diazepam binding inhibitor modulates GABA-mediated inhibition in the brain.

## Introduction

Since their discovery over fifty-five years ago, benzodiazepines (BZDs) have been some of the most widely prescribed drugs in the world, and have remained on the World Health Organization’s list of essential drugs since the list’s inception in 1977 [1]. BZDs are used to treat a variety of conditions, ranging from epilepsy to insomnia to anxiety [2-4], and mediate their effects by binding to the main inhibitory neurotransmitter receptor in the brain, the gamma-aminobutyric acid-A receptor (GABAR) [5]. BZDs modulate GABAR activity, and thus alter neuronal signaling.

GABARs are heteropentamers comprised from nineteen possible subunits (α1-6, β1-3, γ1-3, δ, ε, θ, π and ρ1-3) that are expressed in distinct brain regions, with distinct pharmacological properties. There are two GABA binding sites in the extracellular interfaces between β and α subunits, while BZDs bind in an extracellular pocket at the α/γ subunit interface. BZDs, such as diazepam or flurazepam (FZM), can act as positive allosteric modulators (PAMs), which increase GABA-mediated currents, or negative allosteric modulators (NAMs) like methyl-6,7-dimethoxy-4-ethyl-beta-carboline-3-carboxylate (DMCM) which decrease GABA-elicited currents. Zero-modulators, such as flumazenil (FLZ, Ro15-1788) when acting on α1 and α5-containing GABARs [6], occupy the BZD site but do not alter GABA-elicited currents. The effects of some BZDs like Ro15-4513 depend on GABAR subunit composition. Ro15-4513 acts as PAM when interacting with α_4_β_2_γ_2_ and α_6_β_2_γ_2_ GABARs, but is a NAM when bound to α_1_β_2_γ_2_ receptors [7].

The presence of a binding site on the GABAR for synthetically manufactured BZDs has long suggested the existence of an endogenous synthesized molecule that binds to the same site. In 1983, a candidate peptide was isolated from rat brain homogenates, which displaced tritiated diazepam in a radioligand binding assay [8]. This 10kD, 87 amino-acid protein was named the diazepam binding inhibitor (DBI). Recent DBI knock-down and over-expression studies in the thalamic reticular nucleus demonstrate that DBI works *in vivo* as a PAM where it potentiates GABA-mediated currents and suppresses epileptic activity[9]. In contrast, DBI knock-down in the subventricular zone of the lateral ventricles and in the hippocampal subgranular zone demonstrate that DBI and one of its peptide fragments, ODN, inhibit GABA-induced currents [10, 11], indicating that in these brain regions DBI and ODN work as NAMs of GABARs. We hypothesized that the differences in DBI’s modulatory effects are due to regional differences in GABAR subunit expression and GABAR subunit composition.

Here, we used two-electrode voltage clamping and measured DBI effects on GABA-activated currents from GABARs expressed in *Xenopus laevis* oocytes. Given DBI is highly expressed in the hippocampus and thalamus [12, 13] and published work using DBI examined its effects on inhibition in the hippocampus and thalamic reticular nucleus [9, 11, 14], we used α_5_β_3_γ_2L_ and α_3_β_3_γ_2L_ GABAR subtypes which are particularly enriched in these regions, respectively [15]. We found that DBI positively modulates α_5_β_3_γ_2L_ receptors, whereas DBI is a weak negative modulator of α_3_β_3_γ_2L_ GABARs. We used *in silico* protein-protein docking to visualize potential DBI-GABAR interactions.

## Materials and Methods

### DBI expression and purification for functional testing

We used cDNA for human DBI isoform3 (GeneBank ID CAG33237.1) with a C-terminal histidine tag in a pET28a vector. The plasmid was chemically transformed into BL21 DE3 competent cells, plated on LB-Kanamycin (Kan) agar plates, and grown overnight at 37°C. A single colony was picked and grown overnight in a 5mL culture of LB with 250μg Kan while shaking at 37°C. This culture was used to inoculate 500mL LB + Kan and the culture was grown until OD_600_ reached 0.6-0.8. Protein expression was induced with IPTG (500μM) for 4h at 37°C. After induction, cells were pelleted by spinning at 10,000xg for 10min at 4°C. The bacterial cell pellet was frozen at −80°C. Cell pellet was thawed and resuspended in 50mL Buffer A (sodium phosphate, 50mM, pH 7.4; NaCl, 300mM) and 25mM imidazole (IDA) with a protease inhibitor cocktail (pepstatin, aprotinin, leupeptin). After lysis using an Emulsiflex (ATA Scientific, Taren Point, NSW Australia), samples were spun at 16800xg for 20min, 4°C. The supernatant was bound to an Ni-NTA column which was pre-equilibrated in column buffer (Buffer A and 10mM imidazole, pH 7.4). His-tagged DBI was eluted using elution buffer (Buffer A and 250mM imidazole). A PD10 desalting column was used to remove imidazole, DBI was eluted in Buffer A. DBI molecular weight was verified via 15% SDS-PAGE stained with Coomasie blue and protein level quantified using a Nanodrop 2000 (Thermo Fisher Scientific, Waltham, MA), using an absorption coefficient of 1.68. DBI with the histidine tag attached has a mass of 10.87kDa. All data shown used purified His-tagged DBI.

### DBI purification for NMR

DBI was transformed into BL21 DE3 cells, grown in _15_N-labeled M9 media, and purified as described above. A _1_H-_15_N HSQC spectrum was obtained using a Bruker Avance III 600 MHz spectrometer, with 5-10% D_2_O added to the protein sample. The spectrum was collected at 25°C. Data were processed using NMRPipe [16] and analyzed using CcpNmr analysis [17].

### Expression in *Xenopus laevis* oocytes

Rat cDNA encoding the GABAR α3, α5, β3 and γ2L subunits subcloned into the pUNIV vector [18] were used. cRNA was transcribed from NotI-digested cDNA using the mMessage T7 kit (Ambion, Austin, TX). *Xenopus laevis* oocytes were harvested and prepared as described previously [19]. Oocytes were injected with 27-54nL of GABAR subunit cRNA at 10ng/μL α, 10ng/μL β and 100ng/μL γ2L following previously described methods [20], stored at 16°C in ND96 buffer (96mΜ NaCl, 2mΜ KCl, 1mΜ MgCl2, 1.8mΜ CaCl2, and 5mΜ HEPES, pH 7.2) supplemented with 100μg/ml bovine serum albumin and gentamycin and recorded from 2-7 days after injection.

### Two electrode voltage clamp

Electrophysiological recordings were performed as described previously [21]. Oocytes expressing GABARs were held between −40 and −80mV under a two-electrode voltage clamp and continuously perfused with ND96 at 5mL/min in a 200 μL volume chamber. Boroscilicate glass electrodes (0.4-1.0 MΩ, Warner Instruments, Hamden, CT) were filled with 3M KCl. Data were collected at room temperature using two different volatage clamps. One was a GeneClamp 500 (Molecular Devices, Sunnyvale, CA) interfaced to a computer via a Digidata 1200 device (Molecular Devices) with data recorded using Whole Cell Program, version 3.6.7 (J. Dempster, University of Strathclyde, Glasgow, UK). The other was an Oocyte Clamp OC-725A (Warner Instruments) with a Digidata 1440A (Molecular Devices) with data recorded using AxoScope, pCLAMP 10 (Molecular Devices). Data and results obtained from the clamps were indistinguishable.

A stock solution of 1M GABA (Sigma-Aldrich, St Louis, MO) was made in water, stored at −20°C and thawed before use. GABA dilutions for experiments were prepared fresh daily in ND96. A stock solution of 10mM FZM (Sigma/RBI, Natick, MA) in water was diluted in ND96 for working concentrations daily. A stock solution of 10mM DMCM (Sigma/RBI, Natick, MA) was prepared in dimethyl sulfoxide and diluted daily in ND96 for working concentrations in which the final concentration of dimethyl sulfoxide (≤0.1%) did not affect GABAR function. A stock solution of 2mg/mL aprotinin (Prospec, East Brunswick, NJ) was prepared in Buffer A and diluted daily in Buffer A to a final concentration of 50μM. Aprotinin has a mass of 6.5kDa.

### Concentration-response analysis

GABA concentration-response curves were determined as described previously [21]. Seven concentrations of GABA (1μM, 3μM, 10μM, 30μM, 100μM, 1mM and 10mM) were used to determine GABA EC50 values. Concentration-response data were fit using Prism version 9.1 (GraphPad Software Inc., San Diego, CA) to the equation: I = Imax/[1 + (EC50/[A]nH)], in which I is the peak current response to a given GABA concentration, Imax is the maximal amplitude of GABA activated current, EC50 is the GABA concentration that produces the half-maximal response, [A] is test GABA concentration, and nH is the Hill coefficient.

### Drug modulation

Drug modulation was measured at GABA EC_20_ and evaluated as I_GABA+Drug_/I_GABA_, where I_GABA+Drug_ is the GABA-mediated current in the presence of drug and I_GABA_ is the GABA-mediated current in the absence of drug. To measure drug modulation, a 5s application of GABA EC_20_ was applied and followed immediately by a 5s application of GABA EC_20_+Drug. When multiple drugs were applied to a single oocyte, GABA EC_20_ was applied following drug treatment until current amplitudes returned to initial GABA EC_20_ level to ensure complete washout of drugs between different drug treatments.

### Protein-protein docking

We made α_3_β_3_γ_2L_ and α_5_β_3_γ_2L_ GABAR homology models based on the cryoEM structure of the α_1_β_3_γ_2L_ GABAR obtained in the presence of the BZD alprazolam (PDB 6HUO) [22]. The homology models were built on Sybyl (Tripos Inc). The GABAR α3 or α5 sequences were manually threaded on to the cryoEM model of the α_1_β_3_γ_2L_ GABAR based on their sequence alignment with the α1 subunit using the graphical interface of the Sybyl program. After resolving steric clashes, either manually or by using focused minimization, the resulting structures were subject to whole molecule energy minimization using the Tripos force field. Gasteiger-Huckel charges were added to the atoms, and the dielectric function set at 1.0. After Simplex initial minimization, the Powell function in Tripos was used to minimize to a termination gradient of 0.05 kcal/(mol x angstrom). The GABAR model images were developed using PyMOL (Schrödinger, LLC, New York). BZD alprazolam was removed from models prior to docking.

Protein-protein interactions between the α_3_β_3_γ_2L_ and α_5_β_3_γ_2L_ GABAR models and the unliganded human DBI crystal structure (PDB 2FJ9) [23] were identified using ClusPro, a web-based program for computational docking of proteins [24-26]. ClusPro analyzes 70,000 rotations of the ligand and generates up to 30 highly populated clusters of ligand structures docked in similar locations on the receptor with low energy scores. Clusters are ranked by population size rather than an energy score. Docking clusters selected for analysis were based on their proximity to the BZD binding site. We analyzed the hydrogen bonding between the GABAR alpha subunits and DBI and the GABAR gamma subunit and DBI using Pymol’s hydrogen bonding feature.

### Statistical analysis

All data were from at least three different oocytes from at least two different frogs. Data are represented as mean ± SD. Significant differences in drug modulation between subunit compositions were calculated via Kruskal-Wallis test with a Dunn’s multiple comparisons (Prism v9.1, GraphPad Software Inc, San Diego, CA). This test was selected due to the non-normal distribution of data, as evaluated using a D’Agostino and Pearson test which evaluates skewness and kurtosis and generates a P value based on how much these values differ from a Gaussian distribution [27]. Normalized values are used to compare the effects of drug or DBI between α_3_β_3_γ_2L_ and α_5_β_3_γ_2L_ GABARs. Normalized values were also compared to a hypothetical null value of 1 using a one-sample t-test in Prism.

Statistical differences in GABA+drug versus GABA alone current amplitidues elicited from the same oocyte were calculated using a ratio paired t-test of raw, non-normalized values using Prism software rather than a standard paired t-test. For our data, differences between control and treatment is not a consistent measure of effect. The differences are larger when the control current amplitudes are larger. Thus, the ratio (treated/control) is a more consistent way to quantify the effect of the treatment [28].

The effects of DBI on α_5_β_3_γ_2L_ GABA-elicited currents were more variable than effects observed with FZM, a BZD (coefficient of variation for FZM α_5_β_3_γ_2L_ is 17.2%, DBI is 36.0%). The variation was not correlated with DBI purification batch nor oocyte/frog. We hypothesize that flexibility and conformation dynamics of DBI (11kDa) may contribute to the increased variability of its effects as compared to a small BZD drug.

### Ethics Statement

This study was carried out in strict accordance with the recommendations in the Guide for the Care and Use of Laboratory Animals of the National Institutes of Health. The protocol was approved by the Institutional Animal Care and Use Committee of the University of Wisconsin-Madison (Protocol Number: M005155-R02).

## Results

In order to study the effects of DBI on the GABAR receptor, we needed milligram quantities of pure, non-aggregated DBI folded predominantly in a single conformation. We used a histidine-tagged human DBI construct (His-DBI) inserted into a pET28A vector, which is optimized for inducible expression of protein in bacterial cells [29]. We expressed the His-DBI in BL21 *E. coli* cells and purified the protein as described in Methods. DBI purity and size was evaluated using 15% SDS-PAGE (Fig 1A). Following purification, a single coomassie blue stained band was observed with a molecular weight of approximately 11kDa. DBI with the histidine tag attached has a mass of 10.87kDa. Heteronuclear single quantum correlation (HSQC) nuclear magnetic resonance (NMR) of _15_N-labeled DBI demonstrated that our DBI purification process yielded pure, non-aggregated and folded protein in a single conformation, as the number of peaks were very close to the total number of residues in the His-tagged DBI, spectral dispersion was good, and the linewidths were relatively uniform (Fig 1B). Resolved crystal structures of unliganded DBI purified from human liver (PDB 2FJ9 [23]) and *Chaetophractus villosus* Harderian gland (PDB 2FDQ [30]), as well as a resolved NMR structure of unliganded DBI purified from bovine liver (PDB 2ABD [31]) are also found in a single conformation.

**Fig 1.**
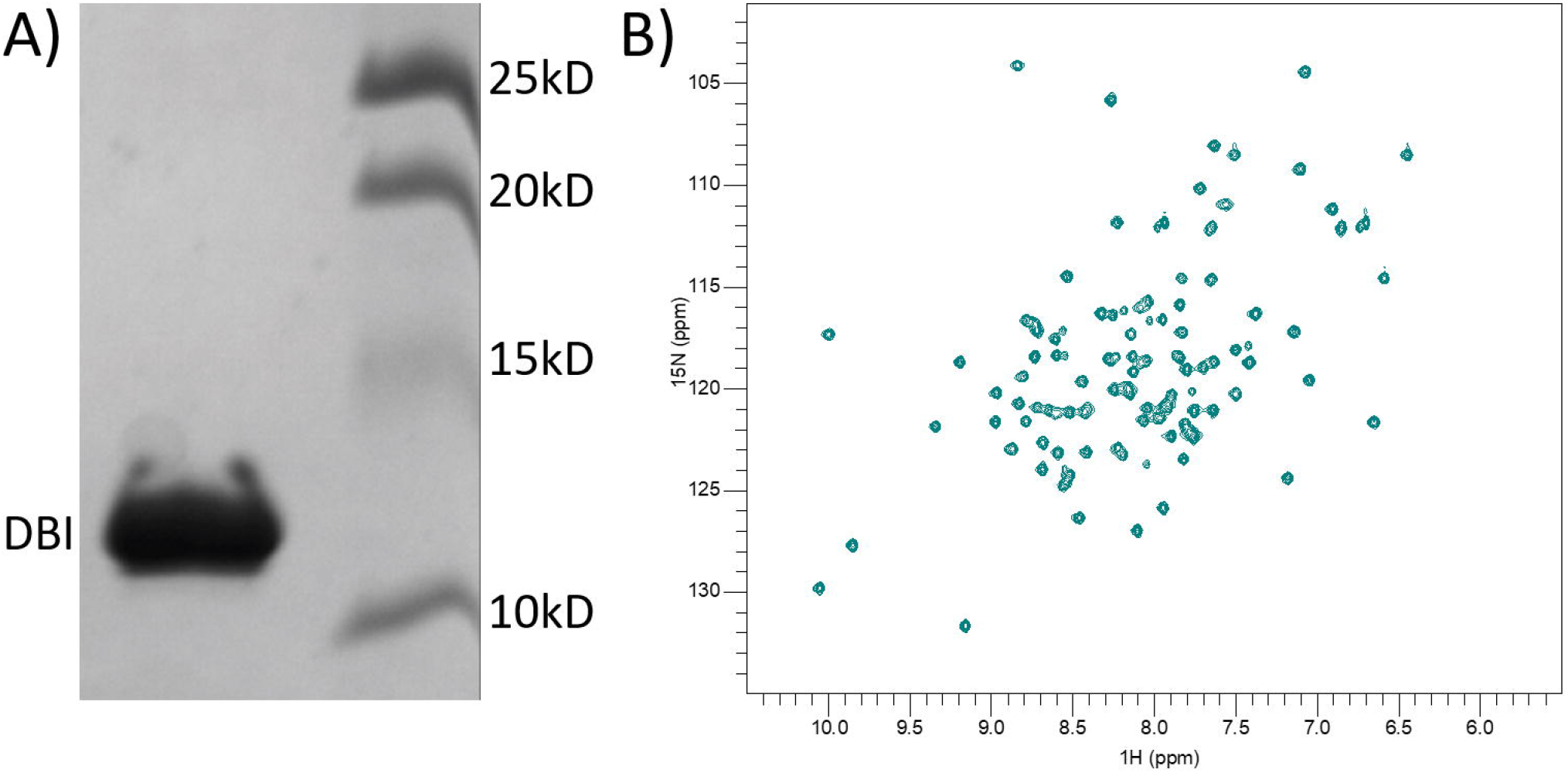
Purified DBI is non-aggregated and folded in a single conformation. (A) 15% SDS-PAGE showing Coomassie blue stained molecular weight marker proteins at 25, 20, 15 and 10kDa, and purified histidine-tagged DBI running at its expected size 10.8kD. (B) _1_H-_15_N HSQC spectrum from DBI grown in _15_N M9 media shows close to expected number of peaks (93 expected, 96 detected), good spectral dispersion and low noise, indicating purified DBI is folded and non-aggregated.

We expressed α_3_β_3_γ_2L_ and α_5_β_3_γ_2L_ GABARs in *Xenopus laevis* oocytes and used two-electrode voltage-clamping to measure and compare the effects of a BZD PAM, flurazepam (FZM), a BZD NAM (DMCM) and DBI on GABA-elicited currents. Initially, we measured GABA concentration responses from oocytes expressing α_3_β_3_γ_2L_ and α_5_β_3_γ_2L_ GABARs. GABA EC_50_ values were 47±11.8 μM, n=5 for α_3_β_3_γ_2L_ GABARs (Fig 2A) and 34±8.6 µM, n=3 for α_5_β_3_γ_2L_ GABARs (Fig 2B), which are consistent with previously published results [32, 33].

**Fig 2.**
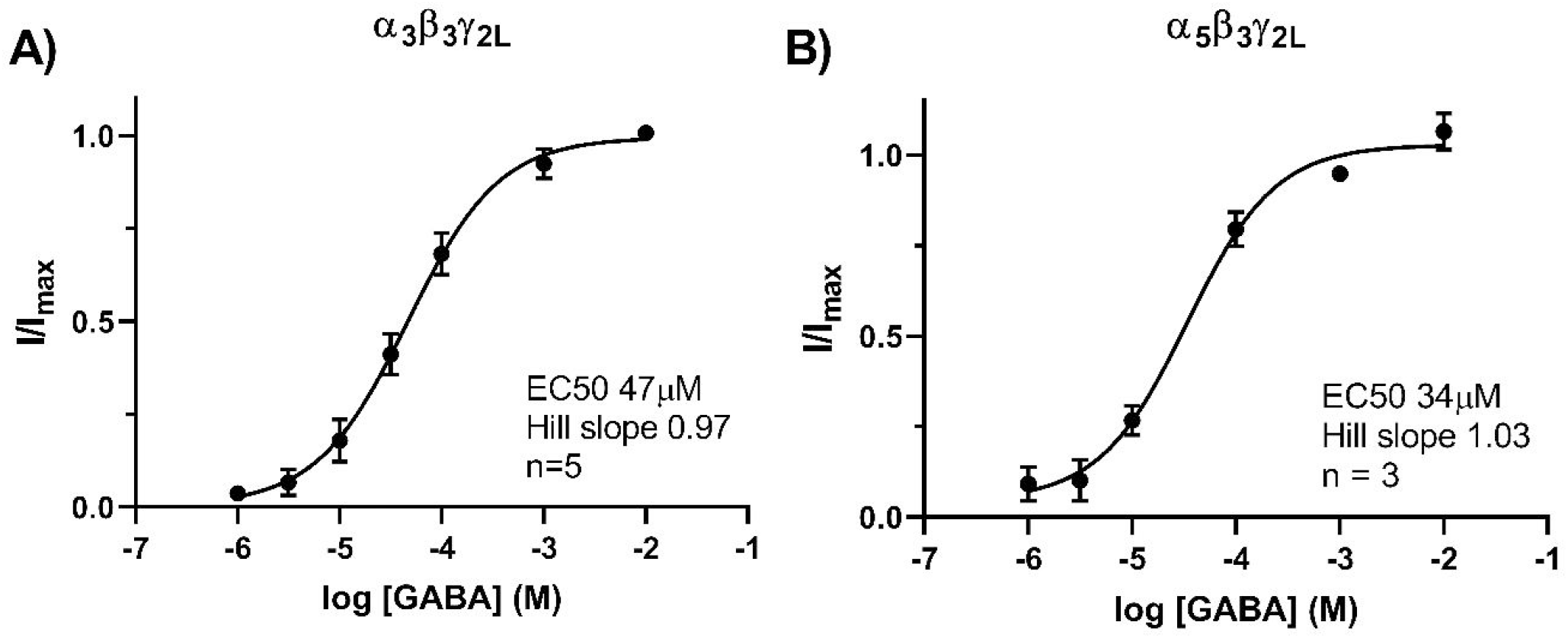
GABA dose response curves for α_3_β_3_γ_2L_ and α_5_β_3_γ_2L_ GABARs. Current responses from oocytes expressing α_3_β_3_γ_2L_ (A) and α_5_β_3_γ_2L_ (B) GABARs were fit by non-linear regression as described in Methods. Data were normalized to maximal GABA current obtained from curve fits and averaged from three to five different oocytes, with error bars showing SD.

To evaluate and compare the effects of the positive BZD modulator, FZM, the negative BZD modulator, DMCM and DBI, we applied GABA EC20 concentration and then co-applied FZM (10μM), DMCM (10μM) or DBI (50μM) with GABA EC20 (Fig 3A). FZM and DMCM concentrations are saturating [34, 35], and the concentration of DBI was selected to reflect the level of DBI and cleavage products measured in rat brains [36]. FZM (10μM) significantly increased GABA currents from oocytes expressing α_3_β_3_γ_2L_ and α_5_β_3_γ_2L_ GABARs 2.7- and 2.0 fold, respectively (Fig. 3; ratio paired t-test p<0.0001 for both GABAR subtypes). The effects of FZM on α3-containing receptors compared to α5-containing receptors were not significantly different (Fig 3B, Dunn’s p>0.999). DMCM (10μM) significantly inhibited GABA currents from both α_3_β_3_γ_2L_ and α_5_β_3_γ_2L_ GABARs (Fig 3, ratio paired t-test p≤0.0001) and the effects of DMCM on α3-containing versus α5-containing GABARs were not significantly different (Fig 3B, Dunn’s p>0.999).

**Fig 3.**
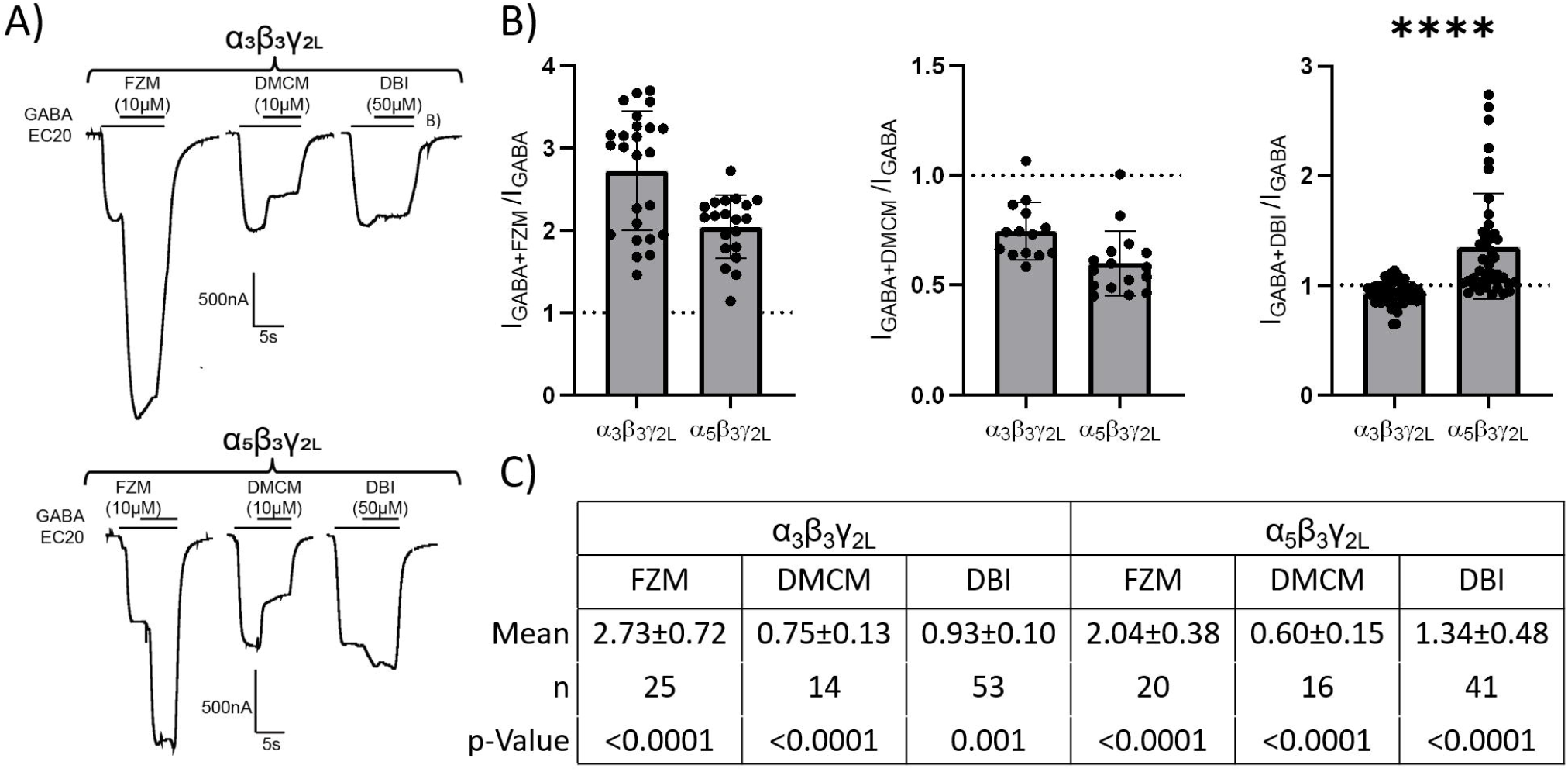
DBI modulates α_3_β_3_γ_2L_ differently than α_5_β_3_γ_2L_ GABARs. (A) Sample current traces from oocytes expressing α_3_β_3_γ_2L_ (top) and α_5_β_3_γ_2L_ (bottom) GABARs showing co-application of FZM (10µM) with GABA EC20 increased GABA current and co-application of DMCM (10µM) decreased current. DBI (50µM) slightly decreased currents from α_3_β_3_γ_2L_ GABARs and increased currents from α_5_β_3_γ_2L_ GABARs. (B) Effects of FZM, DMCM and DBI are plotted as I_Drug+GABA_/I_GABA_. Data from individual occytes are shown as filled circles, bar graphs represent mean +/-SD. Dotted line at 1 represents no effect of drug. DBI effects on α_3_β_3_γ_2L_ and α_5_β_3_γ_2L_ GABARs are significantly different from each other, while the effects of FZM and DMCM are similar for both GABAR subtypes (Kruskal-Wallis test). (C) Table displaying mean +/-SD of FZM, DMCM and DBI effects on currents from α_3_β_3_γ_2L_ and α_5_β_3_γ_2L_ GABAR, n = number of oocytes. Currents in the presence of drug/DBI are all significantly different than GABA currents alone (ratio paired t-test p≤0.001).

We used the same methods to evaluate the effects of purified DBI (50μM) on GABA EC20 currents from α_3_β_3_γ_2L_ and α_5_β_3_γ_2L_ GABARs. DBI weakly inhibited GABA currents from α_3_β_3_γ_2L_ GABARs (Fig. 3). Current amplitudes in the presence of DBI were significantly less than from currents elicited by GABA alone (Fig. 3C, ratio paired t-test p=0.001). In contrast, DBI significantly potentiated GABA currents from α_5_β_3_γ_2L_ GABARs 1.3-fold (Fig 3, ratio paired t-test p<0.0001). The data indicate that DBI acts as a very weak NAM for α3-containing receptors and a PAM for α5-containing receptors, with a significant difference in the effects of DBI based on subunit combination (Dunn’s p<0.0001, Fig 3B).

Since the effect of DBI on α_3_β_3_γ_2L_ GABARs was modest, we measured GABA currents elicited from repeated pulses of EC20 GABA as a control. There were no significant differences in the amplitudes elicited by two EC20 GABA pulses prior to GABA+DBI application (GABA_1_ and GABA_2_, ratio paired t-test α_3_β_3_γ_2L_ p=0.08, α_5_β_3_γ_2L_ p=0.08, S2A-B). Moreover, GABA-elicited current amplitudes recovered to pre-DBI size after DBI was washed off. GABA currents preceeding DBI (GABA_2_) and following GABA+DBI (GABA_3_) were not statistically different from each other (ratio paired t-test α_3_β_3_γ_2L_ p=0.36, α_5_β_3_γ_2L_ p=0.33, S2C-D). Normalized GABA_1_ and GABA_3_ (I_GABA_/I_GABA2_) were not significantly different than 1 (α_3_β_3_γ_2L_ GABA1 p=0.22, GABA3 p=0.97; α_5_β_3_γ_2L_ GABA1 p=0.24, GABA3 p=0.19; S2E-F) whereas normalized GABA+DBI were significantly different (p<0.0001). Thus while the effect of DBI on α_3_β_3_γ_2L_ GABARs was modest, it is not due to a solution artifact.

In order to examine whether the effects of DBI were specific, we measured the effects of aprotinin, a small soluble protein like DBI, on GABA currents elicited by EC20 GABA from α_5_β_3_γ_2L_ GABARs. Co-application of 50μM aprotinin did not affect GABA-elicied amplitudes (ratio paired-test p=0.64, S3), indicating that the effects of DBI are specific and are not due to just applying a small protein to the GABAR.

α3 and α5 are homologous GABAR subunit isoforms with many conserved residues (73% identity) [37]. To explore potential mechanisms underlying DBI’s actions on α3-containing versus α5-containing GABARs, we constructed α_3_β_3_γ_2L_ and α_5_β_3_γ_2L_ GABAR homology models and examined protein-protein interactions with DBI using ClusPro, a web-based program for computational docking of proteins [24-26]. This program generates clusters which represent populations of structures where the ligand has docked with low energy at similar locations on the protein. Clusters are ranked based on population size.

For both α_3_β_3_γ_2L_ and α_5_β_3_γ_2L_ GABARs, ClusPro generated many highly populated clusters with DBI bound at the BZD α/γ binding site interface (Fig.4). A flexible loop between the second and third α helices of DBI penetrates under Loop C of the BZD binding site. Residues in this loop code for a DBI proteolytic peptide call octadecaneuropeptide (ODN) that has been shown to displace _3_H-BZD binding [38]. We selected two clusters, which showed the closest ODN-BZD site interactions, for further analysis. Fig 4A shows DBI docked at GABAR α_3_/γ_2L_ and α_5_/γ_2L_ subunit interfaces. Fig 4B highlights differences in residues in the α3 and α5 GABAR subunits, located at the BZD binding site interface, that are within H-bonding distance to DBI. Fig 4C highlights differences in DBI residues that interact with the α3 and α5 GABAR subunits. Residues in the GABAR γ2L subunit interacting with DBI and residues in DBI interacting with the γ2L subunit are displayed in Figs. 4D and E, respectively.

**Fig 4.**
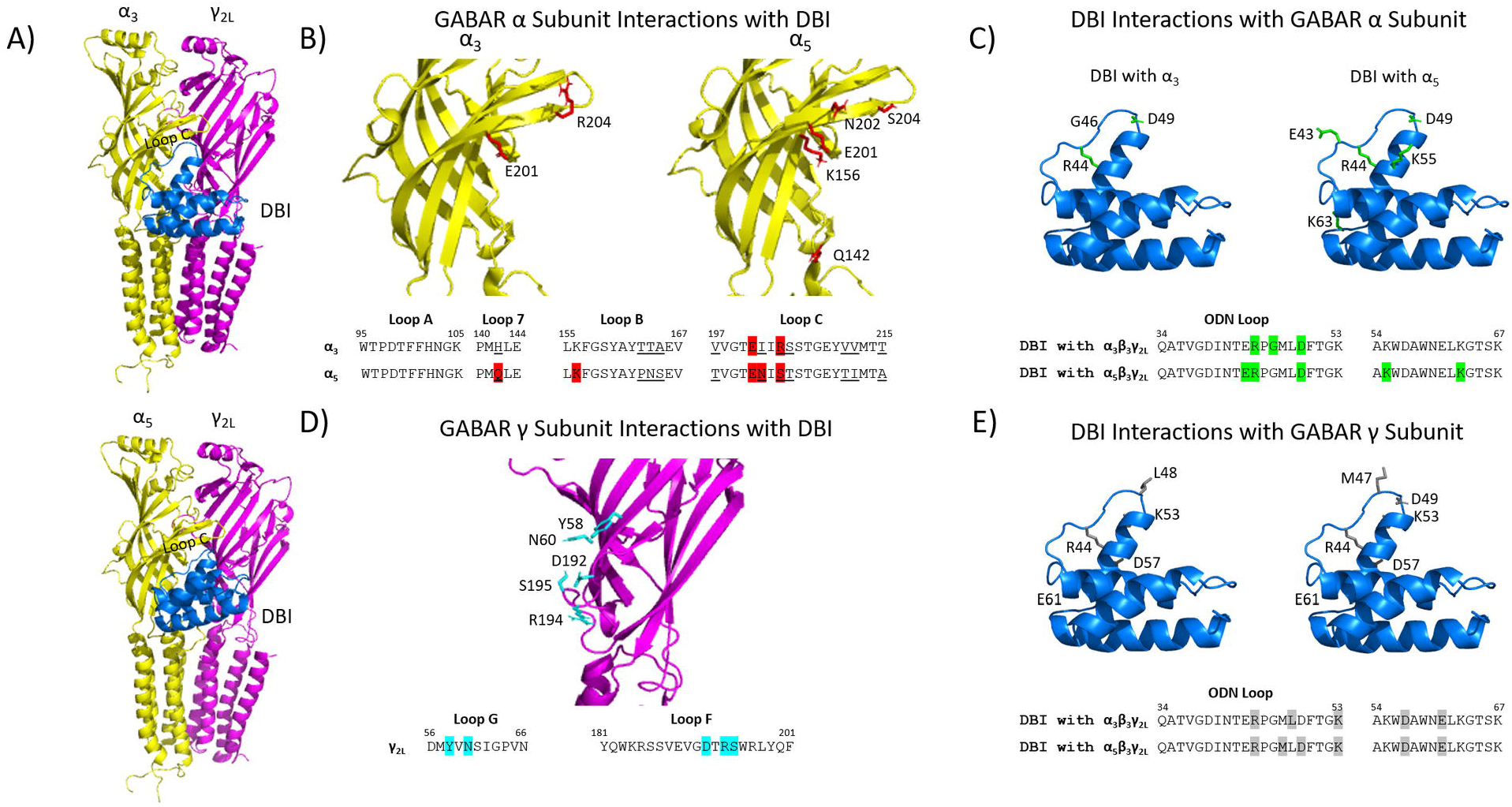
Computational docking reveals DBI can bind at the GABAR α/γ BZD binding site interface. (A) Side views of α_3_/γ_2L_ and α_5_γ_2L_ GABAR subunits (α-yellow, γ-magenta) from homology models based on α_1_β_3_γ_2L_ cryoEM model (PDB 6HUO) showing DBI in blue (PDB 2FJ9) bound at the BZD α/γ binding site interface with flexible ODN loop under Loop C of the BZD binding site. *In silico* dockings performed using ClusPro as described in Methods. B) Extracellular domains of α3 and α5 GABAR subunits with residues predicted to H-bond to DBI shown in stick and colored red. α3 and α5 GABAR sequence alignments shown below, with H-bonding residues highlighted in red and non-identical residues underlined. C) DBI with residues predicted to H-bond to α_3_ and α_5_ subunits shown in stick and colored in green. DBI sequences shown below with H-bonding residues colored green. D) Extracellular domain of γ GABAR subunit with residues predicted to H-bond to DBI shown in stick and colored blue. E) DBI with residues predicted to H-bond to γ subunit shown in stick and colored grey. DBI sequences shown below. For the highlighted hydrophobic residues such as DBI G46, M47, and L48, H-bonding is via the backbone, not the residue side-chain.

When comparing DBI bound to α3 and α5-containing receptors, several differences in interactions between DBI and the GABAR were observed. The GABAR α5 subunit had more residues predicted to H-bond to DBI than the α3 subunit. For example, the α3 histidine at position 142 in Loop7 is not predicted to H-bond with DBI but the aligned α5 glutamine does. In Loop C, the α3 Ile202 does not H-bond with DBI but the aligned α5 asparagine does. In addition, DBI had more H-bond interactions with the γ subunit of α5-containing GABARs. Overall, the computational dockings provide support for the idea that DBI can bind at the BZD binding site interface of GABARs with its flexible ODN loop buried in the BZD binding pocket. The docking is consistent with published work from the Huguenard group showing that adding flumazenil or mutating the BZD site residue α3H126 reduces DBI effects on GABA currents linking DBI’s functional modulation to action at the BZD binding site [9]. The dockings provide a framework for revealing structural mechanisms underlying DBI effects on GABARs.

## Discussion

Despite the fact that DBI was identified as a putative endozepine in the mid-1980s [8], the mechanisms underlying DBI’s actions are still unclear. Due to DBI’s intracellular roles in long-chain fatty acid metabolism, and its multiple biologically active cleavage products [39-42], direct effects of DBI on GABAR function have not been extensively studied. DBI is highly expressed across many tissue types, with high expression in liver, breast, blood, and the brain. In the brain, mRNA DBI expression patterns obtained from the Human Protein Atlas show DBI expression is high in the hippocampus, amygdala, thalamus and midbrain [43, 44].

The use of flumazenil, Ro15-1788 (FLZ, a BZD binding site antagonist for some GABAR subtypes) provides indirect evidence for the existence of an endogenous GABAR modulator. Application of this BZD zero-modulator increases the rate of decay or decreases the amplitude of GABA-evoked inhibitory postsynaptic currents in neurons in the hippocampus [45], dentate gyrus [46], neocortex [47] and the thalamic reticular nucleus [9], suggesting that FLZ is displacing an endogenous GABAR modulator. Recording from cells in the thalamic reticular nucleus suggest that DBI positively modulates GABARs [9] while studies in the subventricular zone and hippocampal subgranular cells suggest DBI is a NAM [10, 11]. In this study, using purified DBI and heterologous expression of GABARs, we found that DBI is a PAM for α5-containing GABARs and a weak NAM for α3-containing receptors demonstrating that effects of DBI on GABAR activity are regulated by receptor subunit composition.

Recordings from cells in the hippocampal subgranular zone suggest that DBI or its peptide cleavage product, ODN, acts as a NAM [11]. Given α5 subunits are highly expressed in the hippocampus [48], we expected that heterologously expressed α5-containing receptors would be negatively modulated by DBI versus positively modulated, which we observed (Fig. 3). Similarly, patch clamp recordings from the thalamic reticular nucleus where α3 is highly expressed [49] suggested that these receptors are positively modulated by DBI [9], while our experiments with heterologously expressed α_3_β_3_γ_2L_ GABARs demonstrated that DBI acts as weak NAM (Fig. 3).

The differences between data reported here and previously reports are likely due to experimental systems used. In neurons, GABARs are likely associated with accessory subunits such as GARLH or Shisa7 [50, 51], which may alter DBI actions. Furthermore, α3 and α5 are not the only α subunits expressed in the thalamic reticular nucleus and hippocampus, respectively. In native cells, the presence of GABARs comprised of other subunits that may be modulated by DBI make it difficult to assign DBI’s effects to one specific GABAR subtype. Here, using a reductionist approach, we demonstrate that DBI is a PAM of α_5_β_3_γ_2L_ GABARs and a weak NAM of α_3_β_3_γ_2L_ GABARs.

α_5_β_3_γ_2L_ GABARs are highly expressed in the hippocampus and can be found both synaptically and extrasynaptically[52]. α5-containing GABARs play a large role in hippocampal-dependent forms of learning and memory [53-56]. Our data demonstrating that DBI acts as a PAM for α5-containing receptors suggests that DBI should have effects on learning and memory. Consistent with this idea, DBI-knockout mice display a disruption in spatial learning and memory [57] and social behavior [58]. While somatic IPSC recordings from hippocampal CA1 and dentate gyrus regions found that the DBI knockout did not change the effects of FLZ on mIPSPs [14], these fast IPSCs recorded from the soma of pyramidal neurons are unlikely to capture the effects of DBI on the α5-containing receptors that are expressed primarily in the distal dendritic regions, which are involved in hippocampal learning and memory [59, 60].

Dysregulation of GABA-mediated signaling is implicated in a wide variety of neurological diseases and disorders, including Alzheimer’s [61, 62], anxiety [4], Parkinson’s [63] and epilepsy [2]. Our findings demonstrating that DBI’s effects are dependent on GABAR subunit composition lay an important foundation for understanding how inhibition in the brain is regulated.

## Supporting Information

**S1 – Original 15% SDS-PAGE of Fig 1A**. Lanes 1 and 10 show Biorad’s Precision Protein Plus All Blue protein standards ranging from 10 kD to 75 kD. Lane 2 shows protein content of the supernatant after emulsion and final high-speed spin. Proteins contained in flow-through, first and second washes are observed in lanes 3, 4 and 5, respectively. Sample was then eluted from the NiNTA column with a single band observed at the expected size of 11kD (lane 6). Lane 7 is the flow-through when the sample is loaded onto the PD10 desalting column. Lanes 8 and 9 are 6 and 3μL of the PD10 elution, which represents the final, purified and desalted DBI. Overall, we observed the purification of a single ∼11 kDa protein, which is the expected size of His-DBI. Boxed area indicates which region of the gel was shown in Fig 1A.

**S2 –GABA current stability and reversibility of DBI effects**. Currents elicited from two sequential pulses of EC20 GABA (GABA_1_, GABA_2_) from the same oocyte are plotted for α_3_β_3_γ_2L_ (A) and α_5_β_3_γ_2L_ (B). Paired GABA_1_ and GABA_2_ currents were not significantly different (α_3_β_3_γ_2L_ ratio paired t-test p=0.08, α_5_β_3_γ_2L_ ratio paired t-test p=0.08). Currents elicited from EC20 GABA before and after GABA+DBI application (GABA_2_, GABA_3_) from the same oocyte are plotted for α_3_β_3_γ_2L_ (C) and α_5_β_3_γ_2L_ (D). Paired GABA_2_ and GABA_**3**_ currents were not significantly different (α_3_β_3_γ_2L_ ratio paired t-test p=0.36, α_5_β_3_γ_2L_ ratio paired t-test p=0.33) demonstrating that the effects of DBI are reversible. Mean normalized values (I_GABA_/I_GABA2_) ± SD are plotted compared to a hypothetical null value of 1 (dotted line) for α_3_β_3_γ_2L_ (E) and α_5_β_3_γ_2L_ (F). One-sample t-tests show no significant effect of GABA_1_ (α_3_β_3_γ_2L_ p=0.22, α_5_β_3_γ_2L_ p=0.24) or GABA_3_ (α_3_β_3_γ_2L_ p=0.97, α_5_β_3_γ_2L_ p=0.19). Effects of GABA+DBI were significant (p<0.0001).

**S3 – Aprotinin does not affect GABA-elicited currents**. A) Sample GABA current traces from α_5_β_3_γ_2L_ GABARs elicited from two sequential applications of EC20 GABA (left) or EC20 GABA immediately followed by EC20 GABA + 50μM aprotinin, a small soluble protein like DBI (right). Co-application of aprotinin had no effect on GABA-elicited current amplitude. B) Plotted are currents from GABA EC20 alone or GABA+Aprotinin normalized to initial GABA EC20 current (I/I_GABA_). Data from individual oocytes are shown as filled circles, bar graphs represent mean +/-SD. Dotted line at 1 represents no effect of drug. (C) Table displaying mean +/-SD of normalized current for GABA or GABA+Aprotinin treatment, n = number of oocytes. Currents elicted by second application of GABA or GABA+Aprotinin were not significantly different than the initial GABA currents (ratio paired t-test GABA p=0.17, aprotinin p=0.64).

**S4 Table – Raw data for GABA dose-response experiments in Fig 2**. Data represent inward GABA-elicted currents in nA.

**S5 Table – Table of raw data from Fig 3**. Data represent inward GABA-elicted currents in nA.

**S6 Table – Table of raw data from Fig S2**. Data represent inward GABA-elicted currents in nA.

**S7 Table – Table of raw data from Fig S3**. Data represent inward GABA-elicted currents in nA.

## Acknowledgments

We thank the lab of Dr. Henzler-Wildman, University of Wisconsin -Madison for their help with obtaining NMR spectra for DBI at the National Magnetic Resonance Facility at Madison.

