## Supplementary figures and images for "The diazepam binding inhibitor’s modulation of the GABA-A receptor is subunit-dependent"

### Supplemental Figure 1

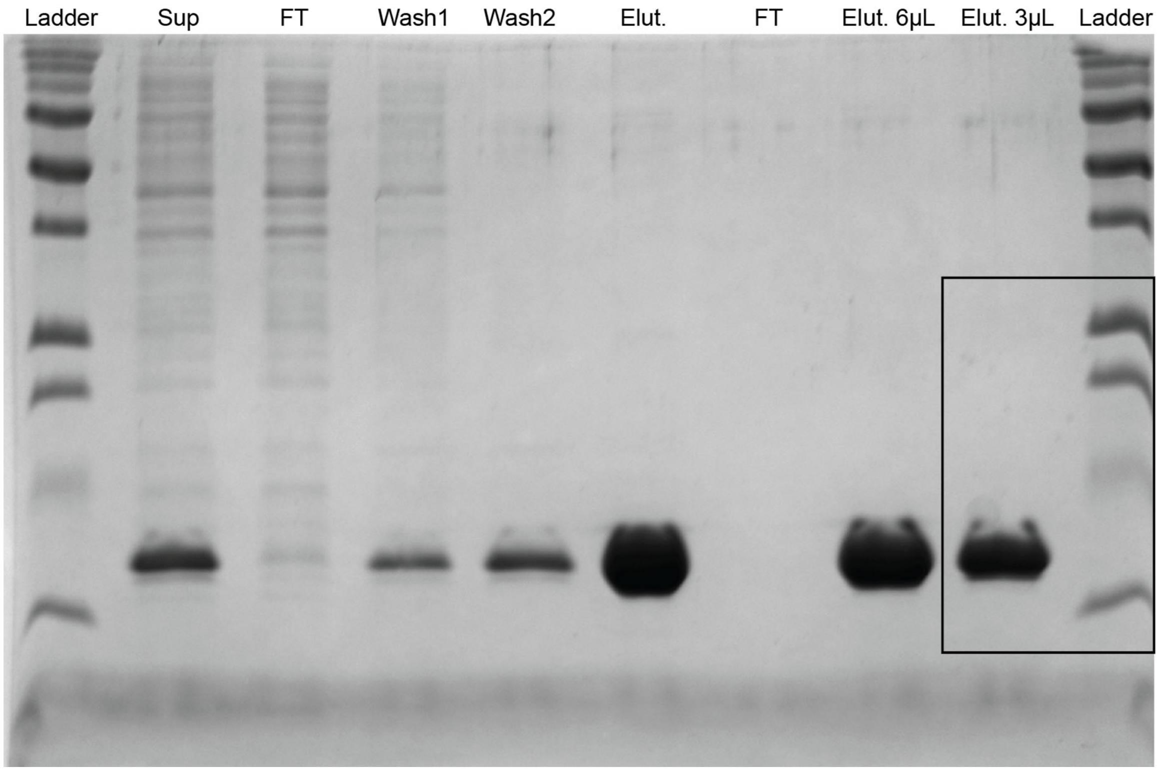

### Supplemental Figure 2

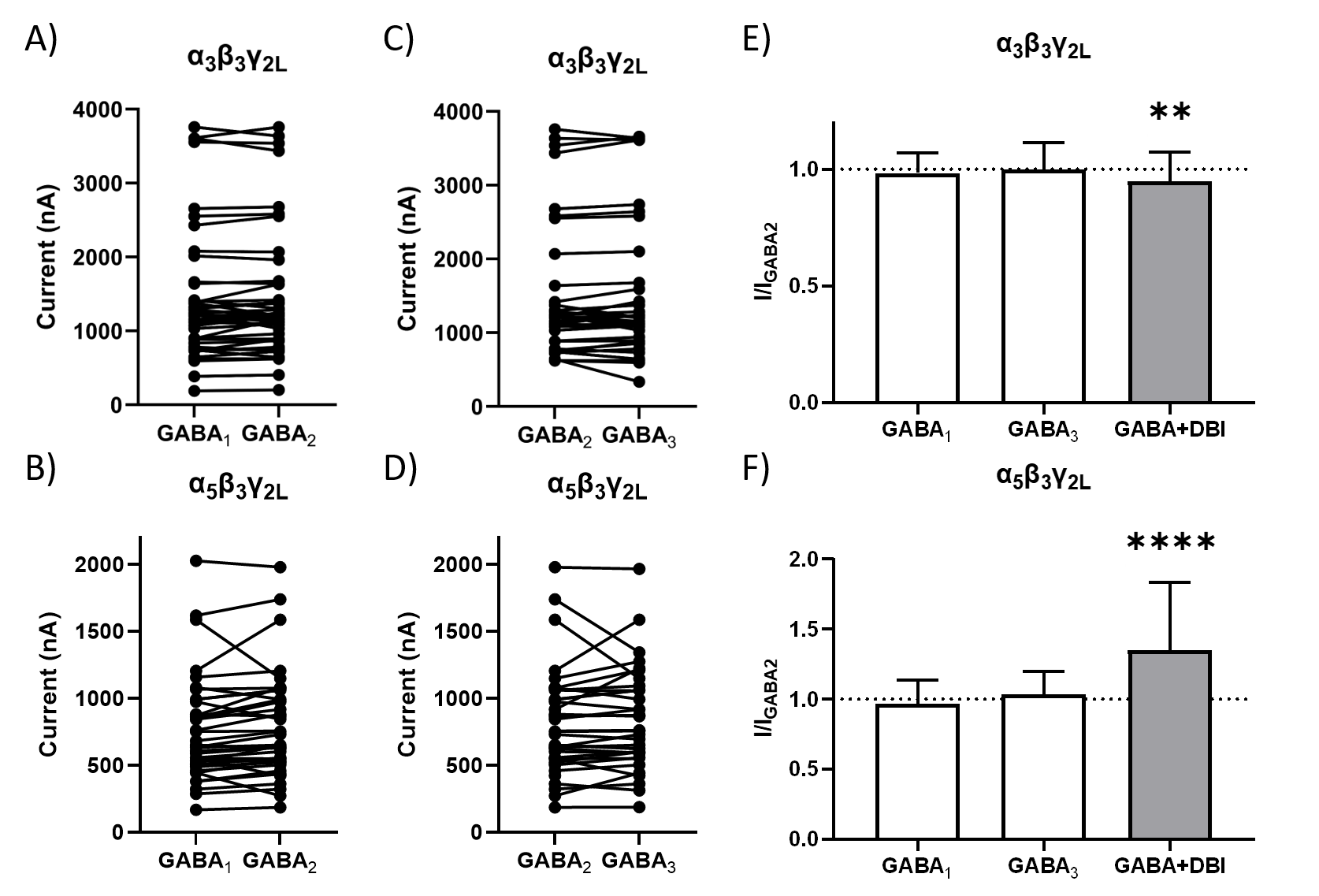

### Supplemental Figure 3

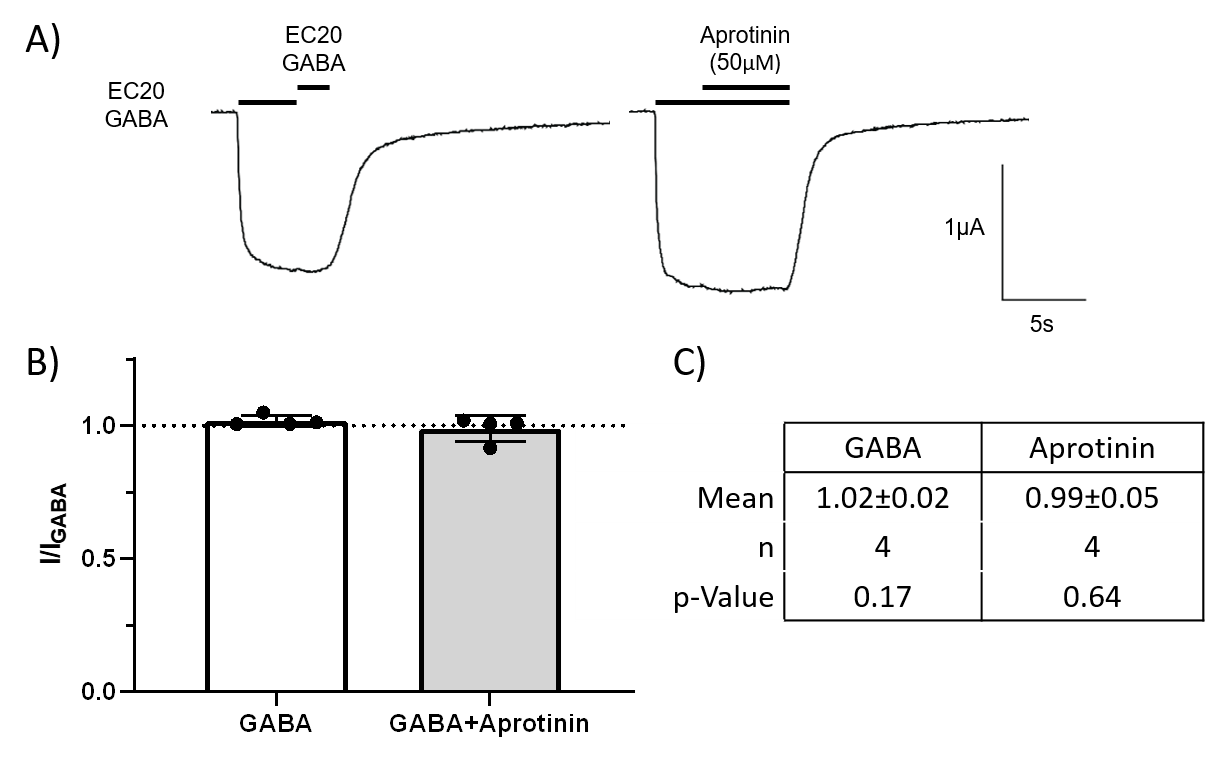
