## Supplemental Table 4 for "The diazepam binding inhibitor’s modulation of the GABA-A receptor is subunit-dependent"

**α3β3γ2L**

| log [GABA] (M) | 9/4/2020 | 9/13/2020 #1 | 9/13/2020 #2 | 9/25/2020 #1 | 9/25/2020 #2 |
|---|---|---|---|---|---|
| -6 | 193.18 | 81.62 | 95.82 | 340.4 | 205.85 |
| -5.5 | 318.16 | 148.15 | 294.01 | 575.54 | 268.5 |
| -5 | 687.9 | 317.8 | 1147.92 | 1768.88 | 906.47 |
| -4.5 | 1917.41 | 546.84 | 2853.23 | 4191.16 | 2329.11 |
| -4 | 3329.65 | 759.19 | 4563.73 | 7651.91 | 4268.66 |
| -3 | 4676.08 | 1009.71 | 5821.81 | 10000 | 6331.81 |
| -2 | 5255.2 | 1147.83 | 6311.04 | 10267.7 | 6856.79 |

**α5β3γ2L**

| log [GABA] M | 11.7.19 #1 | 11.7.19 #2 | 11.17.19 #3 |
|---|---|---|---|
| -6 | 123.08 | 106.88 | 149.45 |
| -5.5 | 139.9 | 151.17 | 119.65 |
| -5 | 265.05 | 455.78 | 509.17 |
| -4 | 693.64 | 1587.79 | 1503.88 |
| -3 | 802.78 | 1794.45 | 1956.39 |
| -2 | 873.43 | 2001.48 | 2282 |
