## Supplemental Table 5 for "The diazepam binding inhibitor’s modulation of the GABA-A receptor is subunit-dependent"

**a3b3g2L**

| GABA | DBI+ GABA | GABA | DMCM+ GABA | GABA | FZM+ GABA |
|---|---|---|---|---|---|
| 202.4 | 229.4 | 191.6 | 170 | 187 | 566.9 |
| 406.1 | 380.3 | 659 | 426.4 | 385.8 | 1308.2 |
| 474 | 399.7 | 943.4 | 604 | 588.8 | 1335.8 |
| 622.5 | 575 | 1189.4 | 694.8 | 645.2 | 2366 |
| 622.8 | 580.2 | 1280.5 | 1109.2 | 735.3 | 2379 |
| 641.8 | 542.6 | 1590.4 | 1030.3 | 841.5 | 2651.2 |
| 725.4 | 727.4 | 1643 | 1043 | 860 | 1793 |
| 732.5 | 727.2 | 1748.9 | 1279.2 | 866.6 | 3088.3 |
| 741.6 | 667.4 | 1832 | 1954 | 894.9 | 2904.6 |
| 762 | 651 | 2710 | 1802.2 | 929.7 | 2738.7 |
| 783 | 859.6 | 2738 | 2034 | 968.5 | 3164.7 |
| 864.9 | 815.8 | 3634.3 | 2768.1 | 1068.7 | 3951.9 |
| 884 | 845.3 | 4019 | 2993 | 1071.4 | 3835.8 |
| 889.1 | 946.3 | 4815.3 | 3991.1 | 1105.6 | 3331.3 |
| 892.9 | 935.2 | | | 1129.7 | 3542 |
| 961.1 | 988.1 | | | 1157.5 | 2192.9 |
| 1032.1 | 1017.8 | | | 1393.9 | 2032.7 |
| 1085 | 1121.1 | | | 1426.3 | 2389 |
| 1090.4 | 1070.2 | | | 1662.6 | 5251 |
| 1094.6 | 988.9 | | | 1710 | 4980 |
| 1117.3 | 1244 | | | 1778 | 3461 |
| 1121 | 1024.2 | | | 2147.9 | 4950.2 |
| 1155 | 1125.9 | | | 2330 | 4532 |
| 1163 | 970.1 | | | 2640.3 | 4963.8 |
| 1205.4 | 1078.7 | | | 3286.4 | 5582 |
| 1206 | 1304.5 | | | | |
| 1206.2 | 1096.7 | | | | |
| 1209.9 | 1170.5 | | | | |
| 1229.4 | 1302 | | | | |
| 1238.7 | 1001.7 | | | | |
| 1240.7 | 1043.6 | | | | |
| 1246.2 | 1076.5 | | | | |
| 1286.3 | 1208.3 | | | | |
| 1301.2 | 1222.6 | | | | |
| 1341.9 | 1296.5 | | | | |
| 1373.5 | 1220.7 | | | | |
| 1384.9 | 1239.7 | | | | |
| 1417.4 | 1395.8 | | | | |
| 1631.3 | 1464.8 | | | | |
| 1637.9 | 1654.9 | | | | |
| 1675.5 | 1516.8 | | | | |
| 1883.2 | 1218.8 | | | | |
| 1962 | 2061.2 | | | | |
| 2068 | 1334.8 | | | | |
| 2551 | 2001 | | | | |
| 2581 | 1948 | | | | |
| 2737.9 | 2463.7 | | | | |
| 3436.4 | 3228.2 | | | | |
| 3634.3 | 3494 | | | | |
| 3657 | 3532.9 | | | | |
| 3757 | 3440 | | | | |
| 4845.3 | 4270.9 | | | | |

**a5b3g2L**

| GABA | DBI+ GABA | GABA | DMCM+ GABA | GABA | FZM+ GABA |
|---|---|---|---|---|---|
| 185.9 | 180.9 | 187.7 | 112.3 | 168.1 | 362.5 |
| 271.7 | 388.5 | 313.3 | 177.7 | 285.7 | 676.6 |
| 321.8 | 880.8 | 468.3 | 470.8 | 380.6 | 870.9 |
| 361.6 | 561.1 | 592.2 | 266.8 | 381.4 | 757.4 |
| 399.7 | 474 | 593.1 | 484.5 | 491.1 | 1069.4 |
| 423.3 | 503.7 | 601.1 | 293.7 | 513.5 | 1397.2 |
| 434.9 | 479.1 | 796 | 362.1 | 525.5 | 1022.5 |
| 441.6 | 487.3 | 873.2 | 508.9 | 533.5 | 1172.3 |
| 460 | 758.2 | 1055.7 | 647.8 | 682 | 1609 |
| 503.8 | 733.8 | 1094.4 | 545.1 | 695.8 | 1660 |
| 521.8 | 590.6 | 1224 | 566.2 | 762.2 | 1623.1 |
| 528.9 | 692.1 | 1342.4 | 877.4 | 841.7 | 1969.6 |
| 549.5 | 561.9 | 1347.9 | 725.9 | 872.4 | 1867.6 |
| 551.7 | 540 | 1590.4 | 1030.3 | 1067.6 | 1898.8 |
| 556.6 | 790.4 | 2120 | 1460 | 1238 | 2855 |
| 602.7 | 1081.3 | 2474 | 1295 | 1297 | 1477 |
| 625.9 | 590.5 | | | 1393.9 | 2032.7 |
| 632.6 | 645 | | | 1618.4 | 2909.6 |
| 635.6 | 943.8 | | | 1697 | 2608 |
| 651 | 604.2 | | | 1981 | 3304 |
| 652 | 661.4 | | | | |
| 729.8 | 1050.8 | | | | |
| 753.1 | 1602.7 | | | | |
| 756.7 | 785.5 | | | | |
| 843.7 | 903.8 | | | | |
| 866.7 | 792.3 | | | | |
| 880.1 | 1088.9 | | | | |
| 918 | 847.9 | | | | |
| 974.2 | 925.4 | | | | |
| 992.6 | 1045.9 | | | | |
| 1060.2 | 1091.2 | | | | |
| 1060.7 | 1051.7 | | | | |
| 1077.9 | 1184.1 | | | | |
| 1080.3 | 1138.9 | | | | |
| 1149.4 | 1183.4 | | | | |
| 1206.2 | 2716.9 | | | | |
| 1501 | 3769 | | | | |
| 1586.3 | 2332.4 | | | | |
| 1739.6 | 2182.6 | | | | |
| 1967 | 2111 | | | | |
| 2286 | 6007 | | | | |
