## Supplemental Table 6 for "The diazepam binding inhibitor’s modulation of the GABA-A receptor is subunit-dependent"

**a3b3g2L**

| $GABA_1$ | $GABA_2$ | GABA +DBI | $GABA_3$ |
|---|---|---|---|
| 187 | 202.4 | 229.4 | |
| 386 | 406.1 | 380.3 | |
| 594 | 622.8 | 580.2 | 622.5 |
| 623 | 622.5 | 575 | 594.8 |
| 645 | 725.4 | 727.4 | 783 |
| 725 | 783 | 859.6 | 733 |
| 732 | 884 | 845.3 | 893 |
| 735.3 | 762 | 651 | 854 |
| 742 | 641.8 | 542.6 | 336 |
| 753 | 741.6 | 667.4 | 642 |
| 783 | 732.5 | 727.2 | 884 |
| 835 | 864.9 | 815.8 | |
| 884 | 892.9 | 935.2 | 889 |
| 893 | 889.1 | 946.3 | 943 |
| 895 | 1163 | 970.1 | |
| 900.1 | 961.1 | 988.1 | |
| 1032 | 1094.6 | 988.9 | 1130 |
| 1069 | 1209.9 | 1170.5 | 1121 |
| 1071 | 1206.2 | 1096.7 | 1229 |
| 1106 | 1246.2 | 1076.5 | 1032 |
| 1117 | 1240.7 | 1043.6 | 1286 |
| 1121 | 1085 | 1121.1 | 1090 |
| 1121 | 1090.4 | 1070.2 | |
| 1155 | 1238.7 | 1001.7 | 1117 |
| 1206 | 1229.4 | 1302 | 1155 |
| 1210 | 1121 | 1024.2 | 1085 |
| 1229 | 1155 | 1125.9 | 1239 |
| 1239 | 1117.3 | 1244 | 1241 |
| 1241 | 1286.3 | 1208.3 | 1205 |
| 1246 | 1032.1 | 1017.8 | 1095 |
| 1286 | 1205.4 | 1078.7 | 1426 |
| 1301 | 1373.5 | 1220.7 | 1206 |
| 1317 | 1384.9 | 1239.7 | |
| 1374 | 1206 | 1304.5 | |
| 1385 | 1631.3 | 1464.8 | |
| 1393.9 | 1417.4 | 1395.8 | 1590.4 |
| 1415 | 1301.2 | 1222.6 | 1374 |
| 1426 | 2699.8 | 1700 | 1415 |
| 1638 | 1675.5 | 1516.8 | |
| 1663 | 1637.9 | 1654.9 | 1678 |
| 2014.2 | 1962 | 2061.2 | |
| 2077 | 2068 | 1334.8 | 2100.5 |
| 2429.5 | 2551 | 2001 | 2581.2 |
| 2550.8 | 2581 | 1948 | 2640.8 |

**a5b3g2L**

| $GABA_1$ | $GABA_2$ | GABA +DBI | $GABA_3$ |
|---|---|---|---|
| 168.1 | 185.9 | 180.9 | 188 |
| 286 | 321.8 | 880.8 | 362 |
| 322 | 361.6 | 561.1 | 313 |
| 381 | 460 | 758.2 | 504 |
| 381 | 434.9 | 479.1 | |
| 442 | 551.7 | 540 | 550 |
| 445 | 271.7 | 388.5 | 442 |
| 460 | 503.8 | 733.8 | 557 |
| 491 | 521.8 | 590.6 | 601 |
| 504 | 556.6 | 790.4 | 603 |
| 514 | 652 | 661.4 | 626 |
| 533.5 | 528.9 | 692.1 | |
| 550 | 423.3 | 503.7 | |
| 552 | 549.5 | 561.9 | 423 |
| 557 | 602.7 | 1081.3 | 593 |
| 589 | 632.6 | 645 | 651 |
| 608.5 | 635.6 | 943.8 | 729.8 |
| 633 | 651 | 604.2 | 622 |
| 636 | 729.8 | 1050.8 | 696 |
| 652 | 625.9 | 590.5 | 592 |
| 682 | 753.1 | 1602.7 | 757 |
| 753 | 756.7 | 785.5 | 762 |
| 762 | 880.1 | 1088.9 | 867 |
| 842 | 974.2 | 925.4 | 918 |
| 844 | 918 | 847.9 | 1224 |
| 860.3 | 991.5 | 1349.4 | 1061 |
| 872.4 | 1080.3 | 1138.9 | 993 |
| 880 | 866.7 | 792.3 | 873 |
| 974 | 843.7 | 903.8 | 918 |
| 992 | 1060.7 | 1051.7 | 1056 |
| 993 | 1060.2 | 1091.2 | 1094 |
| 1067 | 1077.9 | 1184.1 | 1211 |
| 1080 | 992.6 | 1045.9 | 1060 |
| 1158 | 1206.2 | 2716.9 | 1586 |
| 1206 | 1586.3 | 2332.4 | 1149 |
| 1586 | 1149.4 | 1183.4 | 1274.9 |
| 1618 | 1739.6 | 2182.6 | 1342 |
| 2026 | 1979 | 2111 | 1967 |
| | 441.6 | 487.3 | |
| | 1501 | 3769 | |
| | 2286 | 6007 | |

| | | | |
|---|---|---|---|
| 2655 | 2677 | 2463.7 | 2737.9 |
| 3555.8 | 3536.9 | 3532.9 | 3657 |
| 3604 | 3436.4 | 3228.2 | 3613 |
| 3613 | 3757 | 3440 | 3634 |
| 3756.8 | 3634.3 | 3494 | 3613 |
| | 4845.3 | 4270.9 | |
| | 1883.2 | 1883.2 | |
| | 474 | 399.7 | |
| | 1341.9 | 1296.5 | |
| | 474 | 399.7 | |
