## Supplemental Table 7 for "The diazepam binding inhibitor’s modulation of the GABA-A receptor is subunit-dependent"

| GABA | GABA | GABA /GABA |
|---|---|---|
| 1723.3 | 1733.3 | 1.005803 |
| 784.23 | 789.23 | 1.006376 |
| 1173.9 | 1188.8 | 1.012693 |
| 604.41 | 634.38 | 1.049586 |

| GABA | GABA+ Aprotinin | (GABA+ Aprotinin) /GABA |
|---|---|---|
| 1708.3 | 1723.3 | 1.008781 |
| 864.16 | 871.15 | 1.008089 |
| 1543 | 1573 | 1.019443 |
| 889.13 | 814.2 | 0.915727 |
